# Theta-Gamma Cascades and Running Speed

**DOI:** 10.1101/422246

**Authors:** A. Sheremet, J.P. Kennedy, Y. Qin, Y. Zhou, S.D. Lovett, S.N. Burke, A. P. Maurer

## Abstract

The local field potentials (LFPs) of the hippocampus are primarily generated by the spatiotemporal accretion of electrical currents via activated synapses. Oscillations in the hippocampal LFP at theta and gamma frequencies are prominent during awake-behavior and have demonstrated several behavioral correlates. In particular, both oscillations have been observed to increase in amplitude and frequency as a function of running velocity. Previous investigations, however, have examined the relationship between velocity and each of these oscillation bands separately. Based on energy cascade models where “…*perturbations of slow frequencies cause a cascade of energy dissipation at all frequency scales*” (Buzsaki 2006), we hypothesized that the cross-frequency interactions between theta and gamma should increase as a function of velocity. We examined these relationships across multiple layers of the CA1 subregion and found a reliable correlation between the power of theta and the power of gamma, indicative of an amplitude-amplitude relationship. Moreover, there was an increase in the coherence between the power of gamma and the phase of theta, demonstrating increased phase-amplitude coupling with velocity. Finally, at higher velocities, phase entrainment between theta and gamma becomes stronger. These results have important implications and provide new insights regarding how theta and gamma are integrated for neuronal circuit dynamics, with coupling strength determined by the excitatory drive within the hippocampus.

## Introduction

The observation that changes in local field potential (LFP) oscillations correlate with the behavioral state of an animal has a long history (for review, see Buzsáki 2005).Nevertheless, the mechanisms that generate and organize oscillatory activity are still opaque. For example, it has been proposed that different LFP oscillations and their couplings support different processes and network functions (Belluscio et al. 2012; Bieri et al. 2014; Colgin et al. 2009; Gloveli et al. 2005; Lisman and Jensen 2013; Tort et al. 2009; Tort et al. 2008). However, the neurophysiological mechanisms responsible for integrating neuron spiking and oscillations across to temporal scales to support cognition have yet to be elucidated. The observation that hippocampal neuron firing rate (McNaughton et al., 1983; Maurer et al., 2005), theta power (Sheremet et al., 2016 and others) and gamma power (Ahmed and Mehta 2012; Chen et al. 2011; Kemere et al.2013; Zheng et al. 2015) all increase with faster running speeds suggest that synaptic potentials organization across the macroscopic scale (theta) to the microscopic level of the single units. Specifically, as slow oscillations can entrain a large volume of neurons, they are in a better position to ‘coordinate’ local events (Buzsáki and Draguhn 2004).Gamma, on the mesoscopic level, is an inevitable pattern when fast excitation and inhibition compete (Buzsaki 2006; Buzsáki et al. 2004; Buzsáki and Wang 2012; Wang and Buzsáki 1996). On the microscopic scale, basket cells appear to be critical in gamma generation given their fast time course and perisomatic terminals onto pyramidal cells (Buzsaki 2006; Freund and Buzsáki 1996). Thus, during theta states, large-scale activation provides the input necessary for inhibitory and excitatory processes to compete on the mesoscale level (generating high amplitude gamma). This input is also seen on the microscopic level with interactions between interneurons and pyramidal cells.

Previously, we have demonstrated increases in single unit firing rate with rat-running velocity (Maurer et al. 2005). Furthermore, accompanying the increase in firing rate, we found changes in sequence compression (defined as the ratio of the rate of cell assembly transition to the rat’s actual velocity; see Dragoi and Buzsáki 2006; Skaggs et al. 1996) at faster-running speed (Maurer et al. 2012). With respect to the LFP, these changes are accompanied by an increase in theta power, theta frequency, and the number of theta harmonics(Sheremet et al. 2016). Finally, we revealed that pyramidal cells and interneurons interact through putative monosynaptic connection to organize spike firing times (Maurer et al. 2006). Therefore, based on these findings and the role of interneurons in shaping gamma oscillations, we sought to empirically test the hypothesis that theta and gamma inevitably couple in proportion to the amount of activity into the network.

Here, we investigate the interaction between gamma (50-120 Hz; Bragin et al. 1995; Chrobak and Buzsáki 1998; Leung 1992; Penttonen et al. 1998; Stumpf 1965) and the 6-10 Hz theta oscillation (GREEN and ARDUINI 1954; Jung and Kornmüller 1938; Vanderwolf 1969) across hippocampal CA1 laminae: stratum oriens, pyramidal, radiatum and lacunosum-moleculare. Critically, these different laminae are associated with distinct afferent input. The hippocampus has a highly conserved anatomical organization with afferent input and intrinsic fiber projections terminating in discrete layers. For example, the stratum lacunosum-moleculare region of CA1 neurons receives input from the layer III of the entorhinal cortex (Witter et al. 1988), nucleus reuniens (Wouterlood et al. 1990) and amygdala (Pikkarainen et al. 1999), while the stratum radiatum and oriens receive CA3 Schaeffer collateral and commissural fibers (Hjorth-Simonsen 1973; Ishizuka et al. 1990). Concerning local projections, different populations of interneurons target distinct hippocampal lamina with parvalbumin positive cells synapsing in the stratum pyramidale and oriens while the somatostatin positive cells target the stratum lacunosum-moleculare (Freund and Buzsáki 1996). Together, these data suggest that there should be a laminar sensitivity to theta-gamma oscillatory coupling as a function of movement speed.

## Materials and Methods –

### SUBJECTS AND BEHAVIORAL TRAINING

All behavioral procedures were performed in accordance with the National Institutes of Health guidelines for rodents and with protocols approved by the University of Florida Institutional Animal Care and Use Committee. A total of five 4-10 months old Fisher344-Brown Norway Rats (Taconic) were used in the present study. This was a mixed sex cohort comprised of r530♂, r538♂, r544♀, r547♂, and r695♀ in order to integrate sex a biological variable and begin to alleviate the disparity in research focused exclusively on males (Clayton 2016). In the present study, we had no *a priori* predictions that sex will alter oscillations of the hippocampus as single unit physiology is relatively consistent in female animals across estrous (Tropp et al. 2005). Upon arrival, rats were allowed to acclimate to the colony room for one week. The rats were housed individually and maintained on a 12:12 light/dark cycle. All training sessions and electrophysiological recordings took place during the dark phase of the rats’ light/dark cycle. Training consisted of shaping the rats to traverse a circular track for food reward (45mg, unflavored dustless precision pellets; BioServ, New Jersey; Product #F0021). During this time, their body weight was slowly reduced to 85% to their arrival baseline. Once the rat reliably performed more than one lap per minute, they were implanted with a custom single shank silicon probe from NeuroNexus (Ann Arbor, MI). This probe was designed such that thirty-two recording sites, each with a recording area of 177 μm_2_, were spaced 60 μm apart allowing incremental recording across the hippocampal lamina. In preparation for surgery, the probe was cleaned in a 4% dilution of Contrad detergent (Decon Contrad 70 Liquid Detergent, Fisher Scientific) and then rinsed in distilled water (Vandecasteele et al. 2012).

### SURGICAL PROCEDURES

Surgery and all other animal care and procedures were conducted in accordance with the NIH Guide for the Care and Use of Laboratory Animals and approved by the Institutional Animal Care and Use Committee at the University of Florida. Rats were initially sedated in an induction chamber. Once anesthetized, the rat was transferred to a nose cone. The head was shaved with care taken to avoid the whiskers. The rat was then transferred to the stereotax, gently securing the ear bars and placing the front teeth over the incisor bar. The stereotaxic nose cone was secured, ensuring that the rat was appropriately inhaling the anesthesia. During surgical implantation, the rats were maintained under anesthesia with isoflurane administered at doses ranging from 0.5 to 2.5%. Next, ophthalmic ointment was applied and “tanning shades”, fabricated out of foil, were placed over but not touching the eyes to minimize direct light exposure. Multiple cycles of skin cleaning, using betadine followed by alcohol was applied prior to the first incision from approximately the forehead to just behind the ears. The remaining fascia was blunt dissected away and bone bleeding was mitigated through application of bone wax or cautery. Once the location of bregma was determined, the site of the craniotomy was located and a 3×3mm contour was drilled out, but not completed. This was followed by the placement of 7 anchor screws in the bone as well as a reference over the cerebellum and ground screw placed over the cortex. Once the screws were secured, a thin layer of dental acrylic (Grip Cement Industrial Grade, 675571 (powder) 675572 (solvent); Dentsply Caulk, Milford, DE) was applied taking care to not obscure the craniotomy location. Finally, the craniotomy location was completed, irrigating and managing bleeding as necessary once the bone fragment was removed. Next a portion of the dura was removed, taking care to avoid damaging the vessels and the surface of the neocortex. Small bleeding was managed with saline irrigation and gel foam (sterile absorbable gelatin sponges manufactured by Pharmacia & Upjohn Co, Kalamazoo, MI; a division of Pfizer, NY, NY). The probe implant coordinates targeted the dorsal hippocampus (AP: −3.2 mm, ML: 1.5 relative to bregma, DV: −3.7 to brain surface).

Once the probe was in place, the craniotomy was covered with silastic (Kwik-Sil, World Precision Instruments, Sarasota, FL) and then secured to the anchor screws with dental acrylic. Four copper mesh flaps were placed around the probe providing protection as well as acting as a potential Faraday cage. The wires from the reference and ground screws were soldered to the appropriate pins of the connector. Adjacent regions of the coppermesh flaps were soldered together to ensure their electrical continuity and the ground wire soldered to the coppermesh taking care to isolate the reference from contact with the ground. Once the probe was secured, the rat received 10cc of sterile saline as well as metacam (1.0 mg/kg) subcutaneously (the non-steroidal anti-inflammatory is also known as meloxicam; Boehringer Ingelheim Vetmedica, Inc., St. Joseph, MO). The rat was placed in a cage and monitored constantly until fully recovered. Over the next 7 days, the rat was monitored to ensure recovery and no behavioral anomalies. Metacam was administered the day following surgery as well. Antibiotics (Sulfamethoxazole/TrimethoprimOral Suspension at 200mg/40 mg per 5 mls; Aurobindo Pharma USA, Inc., Dayton, NJ) were administered in the rat mash for an additional 5 days.

### NEUROPHYSIOLOGY

Following recovery from surgery, rats were retrained to run unidirectionally on a circle track (outer diameter: 115 cm, inner diameter: 88 cm), receiving food reward at a single location. For rats 530, 544 and 695, data is only analyzed from the circle track conditions. In order to deal with low velocities from the circle track datasets, additional datasets for rats 538 and 539 from running on figure-8 track (112cm wide x 91cm length) were used. In this task, rats were rewarded on successful spatial alternations. Only datasets in which the rats performed more than 85% of trials correctly were used. The local field potential was recorded on a Tucker-Davis Neurophysiology System (Alachua, FL) at ∼24kHz (PZ2 and RZ2, Tucker-Davis Technologies). The animal’s position was recorded at 30 frames/s (Tucker-Davis). Spatial resolution was less than 0.5 cm/pixel.

### ANALYSES AND STATISTICS

Velocity was calculated as the smoothed derivative of position. The local field potential data was analyzed in Matlab^®^ (MathWorks, Natick, MA, USA) using custom written code as well as code imported from the HOSAtoolbox (Swami et al. 2003). Raw LFP records sampled at 24 kHz (Tucker-Davis system) were low-pass filtered down to 2 kHz and divided into sequences of 1024 time samples (approx. 0.5 s). The analysis of the LFP in the current study was based on standard techniques used for stationary signals (Papoulis and Pillai 2002; Priestley 1981) as previously described in Sheremet et al. (2016). Briefly, we assume that the LFP time series is a stochastic process, stationary in the relevant statistics, and decompose it using the discrete Fourier transform (DFT). The Fourier transform time sequences were reduced to the non-redundant frequency domain of 1 Hz ≤ ω ≤ 1 kHz (where ω are the analyzed frequencies), with a frequency increment of 1 Hz.

Electrode position along the CA1-dentate axis was determined initially via visual inspection of the LFP, followed by traditional current source density analyses (Bragin et al. 1995; Buzsáki et al. 1986; Mitzdorf 1985; Rappelsberger et al. 1981). Shifts in the phase of theta from stratum oriens to the dentate (Buzsáki et al. 1983; Leung 1984; Winson 1978) as well as the regional distribution of currents – triggered on ripples –

### Power-Power Correlations

To investigate the relationship between theta and gamma power, the LFP recordings were down-sampled to 2 kHz and split into 1-s segments. In order to remove epochs with potential noise, an LFP quality control algorithm was applied to remove segments with either 1) too large or too small variance (>3*mean(variance) or <0.1*mean(variance) or 2) with maximum (or minimum) LFP recording has a 10 STD deviation from the mean (maximum(LFP)>mean(LFP)+10*std(LFP) or minimum(LFP)<mean(LFP)-10*std(LFP)). The mean velocity for each segment was computed and only segments with mean velocity larger than 5 cm/s were included for this analysis. For each segment, the power spectrum was estimated with the multitaper method. The first three DPSS sequences were used to get the tapered spectrum estimations. Tapered spectrum estimations were computed with 2000 FFT points and averaging over these spectrum estimations generated the multitaper spectrum. The power of the LFP in a given frequency range is the area under the power spectrum curve within that frequency range. Each observation in the power-power correlation analysis is the power of theta (6-10 Hz) and gamma (50-120 Hz) for a 1-s segment.

### Phase-power Coherency

The LFP recordings were down-sampled to 2000 Hz, split into 1-s segments, and sorted according to their corresponding average velocity into 4 classes [0,5], (5,15], (15,35], and >35 cm/s. If X(t) is a 1-s time-series segment that belongs to a given velocity class, the corresponding power time series *Y*_f1_(t) at frequency f_1_ was obtained by using wavelet transform. The wavelet transform decomposes X(t) into the time-scale domain, and *Y*_f1_ (t) is the modulus square of wavelet transform at the scale whose central frequency is *f*_1_. The Fourier transforms of time series X(t) and *Y*_f1_ (t) are 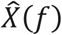 and 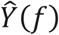 and the cross-spectrum between X(t) and *Y*_f1_ (t) is 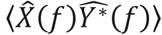 where ⟨. ⟩ is the expected-value operator. The phase-power coherence is defined as the modulus of the cross-spectrum 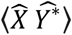 normalized by the power spectra of *X* and *Y*_f1_, and the mathematical expression for phase-power coherence is 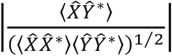 Informally, the phase-power coherence measures how much the power time series of frequency component *f*_1_ is modulated by frequency components *f* in the original signal.

### Bispectrum and Bicoherence

The bispectrum has been thoroughly reviewed in terms of both statistical and mathematical background (Harris 1967) as well as its application to nonlinear wave interaction (Kim and Powers 1979). As noted in our prior publication (Sheremet et al. 2016) as well as others, bispectral analysis (the Fourier transform of the third order cumulant) quantifies the degree of phase coupling between the frequencies of the LFP while the bicoherence quantifies the degree of cross-frequency coupling independent from the amplitude (Barnett et al. 1971; Bullock et al. 1997; Gloveli et al. 2005; Hagihira et al. 2001; Li et al. 2009; Muthuswamy et al. 1999; Ning and Bronzino 1989; Shahbazi Avarvand et al. 2018; Sigl and Chamoun 1994; Wang et al. 2017). The utility of the bispectrum can be illustrated as follows. In the instances of “no coupling”, three independent waves – ω_1_, ω_2_, and ω_1_ + ω_2_, will have statisically independent phases relative to each other, resulting in a random phase mixing when estimated over multiple realizations. In these instances, the bispectrum will take a zero value. When three waves are nonlinearly coupled, however, a phase coherence will exist between ω_1_, ω_2_, and ω_1_ + ω_2_, resulting in a non-zero bispectral value (Fig. 2; Kim et al. 1980). In order to analyze for nonlinear phase coupling between hippocampal LFP frequencies, local field potential the datawere analyzed in Matlab^®^ (MathWorks, Natick, MA, USA) using custom written code as well as code imported from the HOSAtoolbox (Swami et al.2003) for bispectral analysis. It is necessary to reiterate that the nonlinearity of a time-series is expressed by the phase correlation across different frequencies. We used the lowest order (third order) phase coupling, described by the bispectrum and first used in ocean waves by Hasselmann (1963), to analyze the hippocampal LFP, as we previously described in Sheremet et al. (2016). It is emphasized here as well as elsewhere (Pradhan et al. 2012; Van Milligen et al. 1995) that the bispectrum measures phase coupling, defined to occur when the sum of phases between two frequencies is equal to the value of a third frequency plus a constant. In order to associate our analysis with speed, the one-second of LFP for the bispectrum was stored with the mean velocity for the same temporal epoch. Based on mean velocities of every segment, we classified each LFP segment into 4 speed ranges: 0.001 to 5 cm/s; 5 to 15 cm/s; 15 to 35 cm/s; and > 35 cm/s. For statistical comparisons, average nonlinearity was calculated for the 5 to 15 cm/s speed bin andand the > 35 cm/s bin.

**Figure 1:**
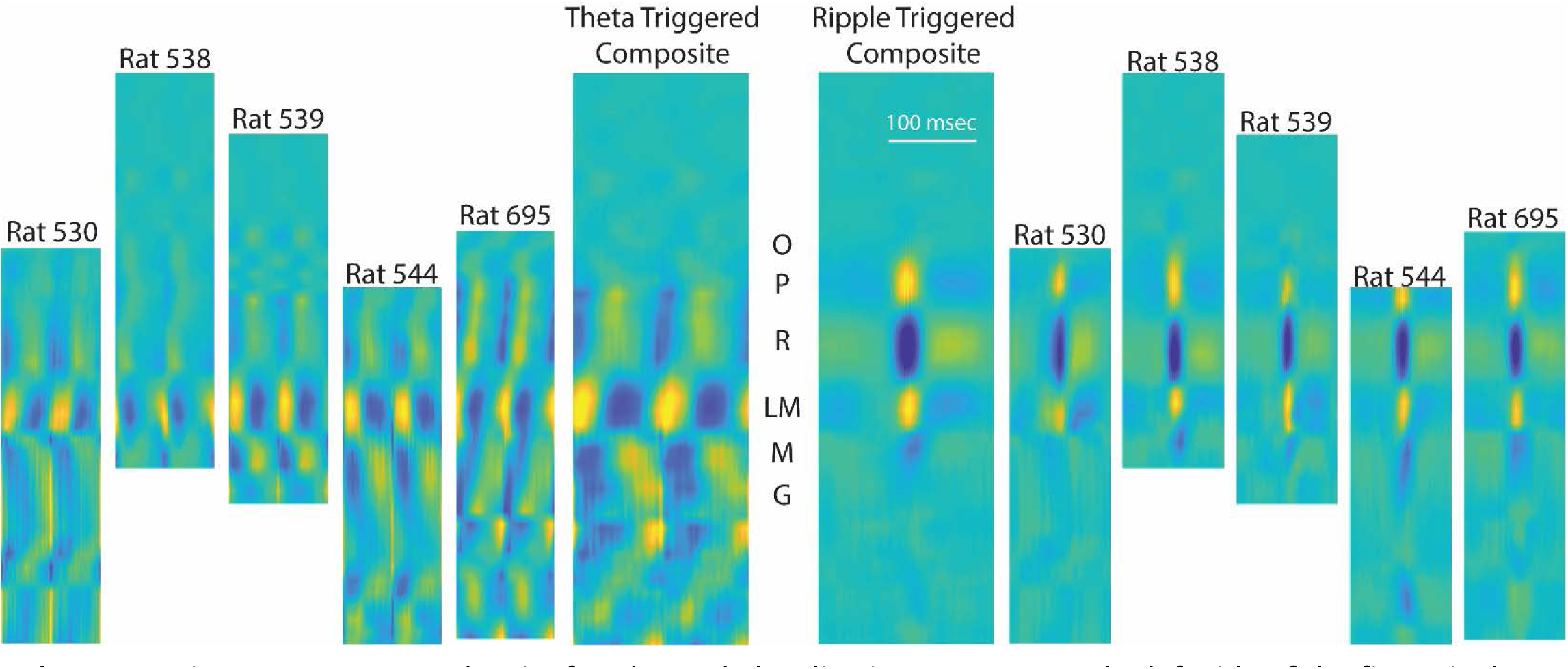
Using current source density for electrode localization across rats. The left side of the figure is the current source density of the raw LFP triggered on the maximum positive deflection of the filtered theta rhythm, with time normalized to two cycles of the hippocampal theta rhythm. On the right are the individual rat current source density plots for the unfiltered LFP, triggered on the maximum positive going ripple in the pyramidal cell layer (sources are warm colors, sinks are cool colors). In order to identify the layers across each rat, composite images (center) were generated with the output compared to prior publications (Bragin et al., 1995; Sullivan et al., 2011; Buzsaki, 2015). Both the theta and ripple current source density analyses were calculated during either awake-behavior or rest epochs. To account for differences in hippocampal size, the current source density images were cross-correlated with each other in order to determine appropriate alignment and compression. For the purposes of the current analyses, the oriens (O), pyramidal layer (P), stratum radiatum (R), lacunosum moleculare (LM), molecular layer (M) and upper granule layer (G) were identified.revealed sources and sinks that directly related to input layers (Fig. 1; Sullivan et al. 2011; Ylinen et al. 1995a).

**Figure 2:**
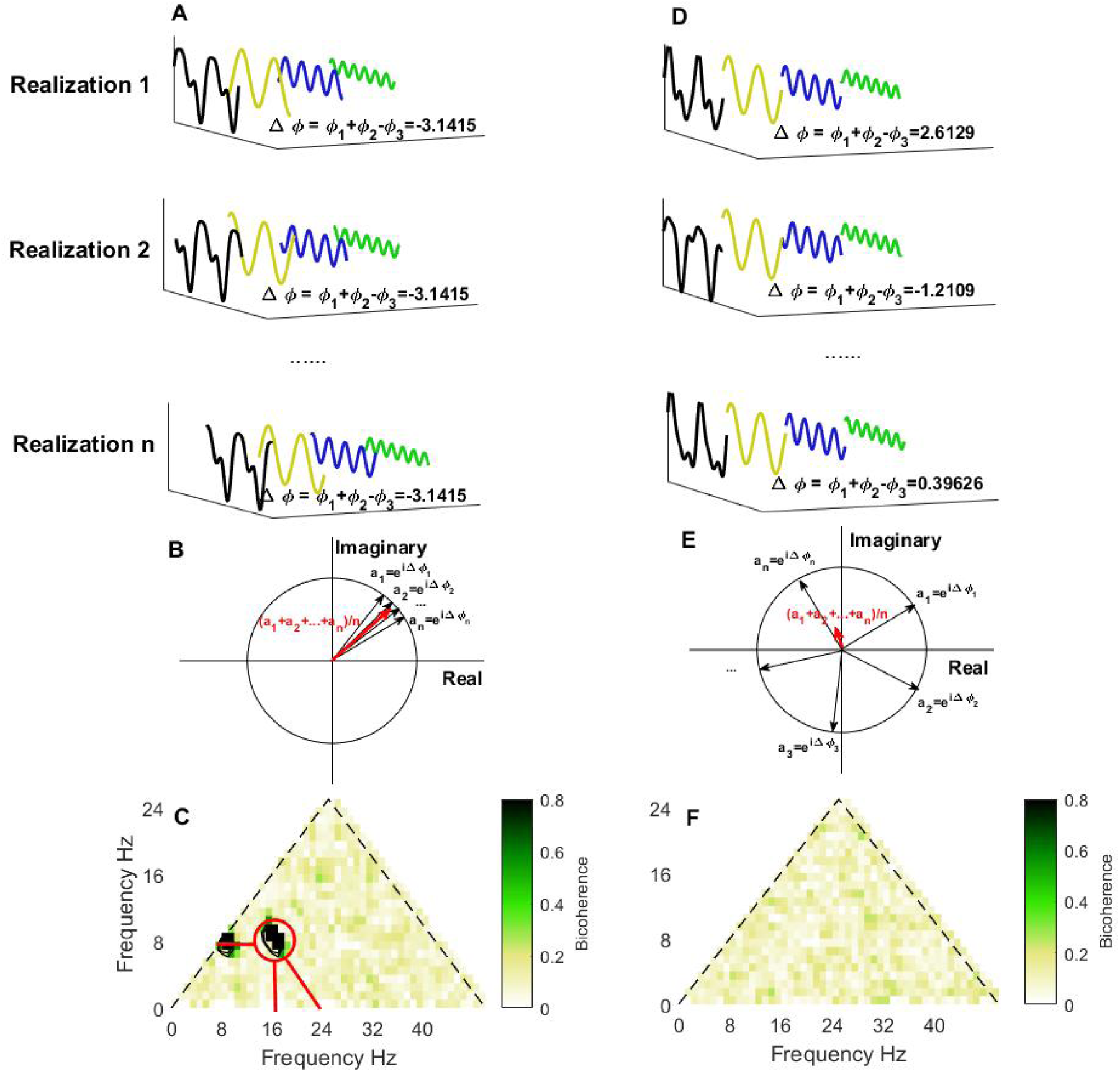
Bicoherence illustration. **A**, Time series with triad phase coupling (black line) resolved into Fourier series where the main components are three sinusoidal waves with frequencies: _*F1=8*_Hz (yellow), *f*_2_ = 16 Hz (blue) and *f*_3_ = *f*_1_ + *f*_2_ = 24 Hz (green). The biphase of the three waves is defined as Δ.ø = ø_*f*1_ + ø_*f*2_ ø_*f*3_ The biphase is almost constant over realizations. Averaging biphase vectors over realizations 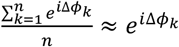 gives a complex number which has the norm. close to 1 as shown in **B**. The frequency triad (f1; f2; f3) is corresponding to the significant region (marked with red circle) in bicoherence plot **C**. Figure **C** shows the modulus of the averaged biphase vectors 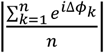 for all the frequency triads. The contour lines in **C** outline the 99% significance level on zero bicoherence which is suggested by Elgar and Guza (1985) as 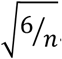 The marked significant region indicates the frequency of three waves: 8 Hz, 16 Hz and 24 Hz (indicated with red lines). The y=xboundary of bicoherence plot **C** is imposed by the symmetry property of bicoherence; and the *x* + *y* = 50 boundary is imposed by the up limit of frequency range of interested _(*f*_*x*_ + *f*_*y*_ = 50 *Hz*). In **C**, there is another significant region (8 Hz, 8Hz & 16Hz) which implies this triad is also phase coupled. **D**, Time series without triad phase coupling (black line) and the Fourier series components with frequency *f*1 = 8 Hz(orange), *f*2 = 16 Hz (blue) and *f*3 = 24 Hz (green). The biphase Δø_ distributes randomly, and the biphase vectors 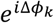 cancel each other when averaging over realizations as shown in **E**, which gives a bicoherence plot without any significant regions in **F**.

## Results

For a preliminary measure of laminar differences of local field potential, we calculated the power spectral density in the CA1 oriens (Or.), stratum pyramidale (CA1.pyr), radiatum (CA1.rad), and lacunosum-moleculare (LM) as a function of speed (**Fig. 3**). Our observations of the power spectra can be characterized in general as having a power law *f* ^−α^ structure. However, unlike previous reports, the spectral slope α is not constant throughout the entire frequency range. Instead, two or more domains with distinct slopes can be identified, most evident in the LM. Using preliminary analyses across layers of the power spectral density, cross-frequency phase-power coherence and bicoherence analyses, these domains formed rather well-defined frequency intervals that can be associated with theta (6-10 Hz) and its harmonics (16Hz, 24Hz, 32Hz, 40Hz and 48Hz; see nonlinear analysis below); gamma (broadly defined here as ∼50-120 Hz similar to Bragin et al. 1995); and fast oscillations (> 120 Hz). Adjacent hippocampal layers also exhibited harmonics of theta, although not as prominent as the LM region. In agreement with previous results (Bragin et al. 1995), theta and gamma power are higher in the LM than the other CA1 layers.

**Figure 3.**
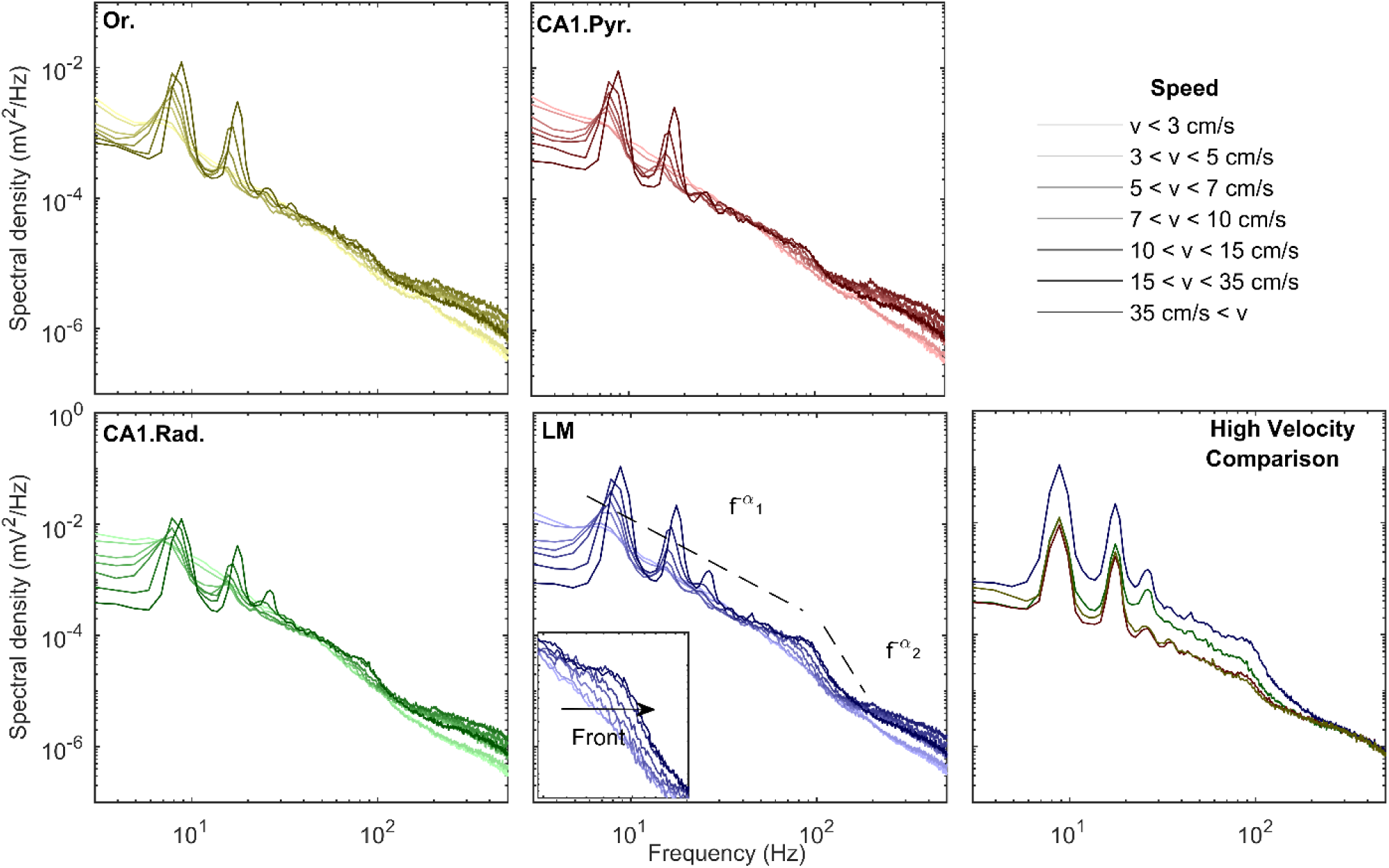
Changes in power spectral density as a function of velocity and CA1 layer (transition from lighter to darker hues indicate low to high velocities). Note that, across all layers, the increases in theta and theta harmonic with speed are associated with an increase in gamma power. Insets show details of the evolution of the gamma range (magnification of gray rectangle area). Spectral shapes in all layers follow a power laws of the type *f*-α. Remarkably, the LM layers exhibits different slopes between the frequency domains that encompass theta and its harmonics and the gamma range. This phenomenon becomes apparent in all regions when comparing all high velocity power spectra across layers (bottom right). Furthermore, as power increases in the gamma range, the spectrum preserves its slope and shifts to the right, creating the appearance of a moving front (inset of LM). Note that, in agreement with prior publications (Bragin et al., 1995), the CA1 oriens and pyramidal layer exhibit the least energetic gamma range of all the layers examined.

In the 50-120 Hz frequency range, the distribution of power acquires the form of a front of constant slope shifting to the right, toward higher frequencies as running speed increases (inset in Figure **3**). This effect is observed across all layers. While the increase in theta power with velocity is in congruence with models by which theta is driven by an external input (Buzsaki 2002; Holsheimer et al. 1982), the shifting gamma front is indicative of a change in circuit dynamics. Specifically, previous work on central pattern generators suggest that high-frequency oscillations can increase as a function of neural drive (Shik et al. 1969). To test this hypothesis relative to the hippocampus, we examined power-power correlations between the theta and gamma bands. If the gamma frequency component of the LFP is excited by the same input as theta, the two oscillatory frequencies should exhibit “power-power” correlations. Therefore, we calculated power for hippocampal theta (narrowly defined as 6-10Hz for this analysis) relative to gamma power (using a range from 50-120Hz) in 1-second windows. By correlating the power of the hippocampal theta rhythm against the power of the gamma within the same region, it is possible to determine whether a power-power relationship exists (**Fig. 4**). Across lamina and rats there was a significant relationship between theta power and gamma power with, on average, 16% of the variance in gamma power being explained by theta power (p = 0.04). Interestingly, the correlation coefficients between theta and gamma power significantly differed between hippocampal lamina (F_[3,12]_ = 10.21, p = 0.001). In fact, the correlation between theta and gamma power was highest in the CA1. Rad (r = 0.51) and lowest in the CA1.Pyr (r = 0.24). Simple orthogonal contrasts comparing the CA1.Rad to the other CA1 lamina indicated that the theta-gamma power correlation was significantly greater in the CA1.Rad compared to the Or (F_[1,4]_ = 19.74, p = 0.01) and the CA1.Pyr (F_[1,4]_ = 16.98, p = 0.02). In contrast, the correlation between theta and gamma power was not significantly different between the CA1.Rad and LM (F_[1,4]_ = 2.11, p = 0.22). These data indicate that theta and gamma power are strongly correlated across all layers, and that this relationship is particularly evident in the CA1.Rad and LM.

**Figure 4.**
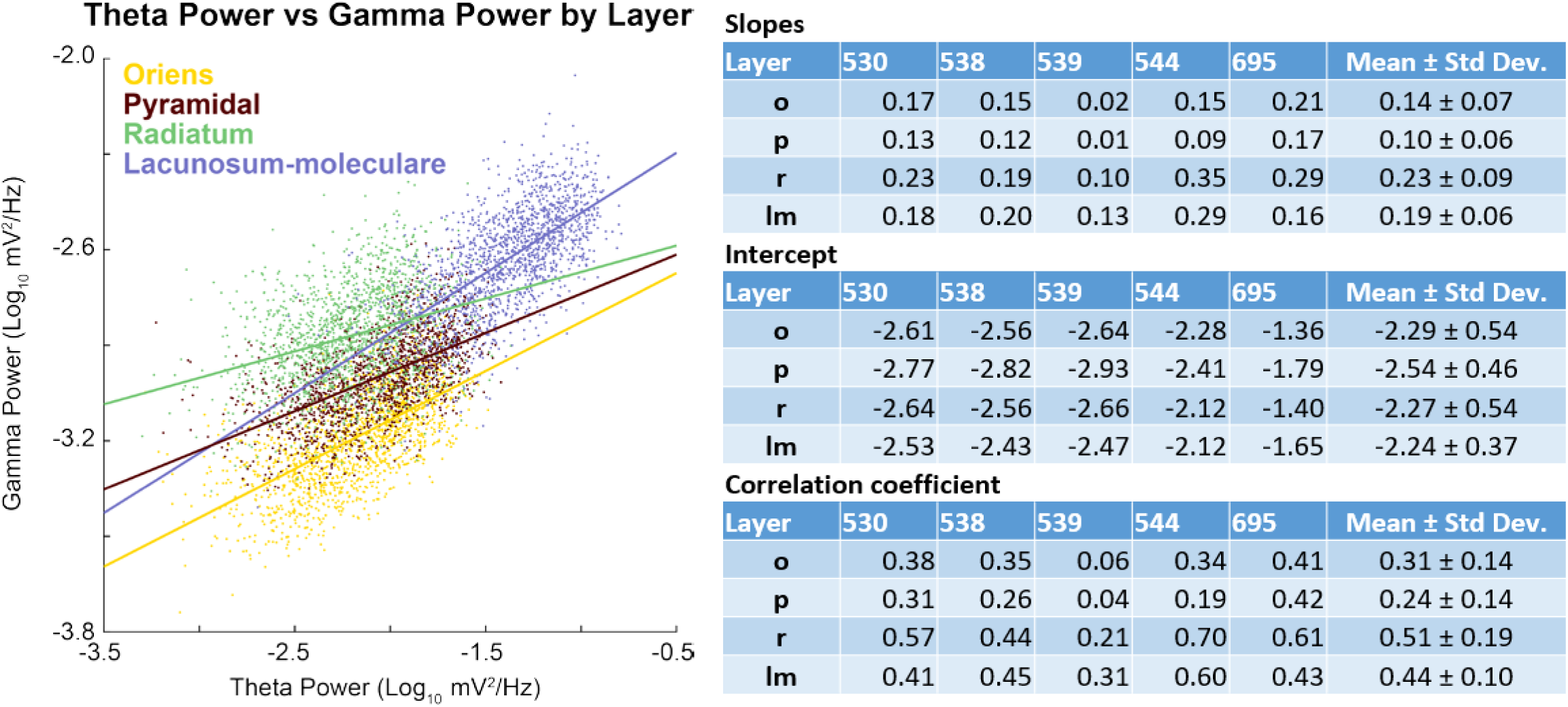
Correlation of theta power to gamma power. (**Left**) Scatter plot and least-squared regression lines for theta and gamma power across the layers of rat 695. Note that there is a clear positive trend across all layers, although strongest in the lacunosum-moleculare. (**Right**) Table containing the individual rat data (slope, intercept and correlation coefficient) by layer as well as averages.

The slope of the relationship between theta and gamma power provides information regarding the rate at which activity transfers across frequency bands. The slope was significantly different across hippocampal laminae (F_[3,12]_ = 7.52, p = 0.004). The slope was greatest in the CA1.Rad (b = 0.23 where b stands for the slope) and smallest in the CA1.Pyr (b = 0.11). Simple orthogonal contrasts comparing the CA1.Rad to the other CA1 lamina indicated that the slope of the theta-gamma power relationship was significantly greater in the CA1.Rad compared to the Or (F_[1,4]_ = 10.64, p = 0.03) and CA1.Pyr (F_[1,4]_ = 14.63, p = 0.02). In contrast, the slope of the theta-gamma power relationship was not significantly different between the CA1.Rad and LM (F_[1,4]_ = 2.11, p = 0.22). This observation is not trivial as it runs counter to contemporary models in which neural oscillations considered to be independent and multiplexed (Akam and Kullmann 2014; McLelland and VanRullen 2016). Multiplexing implies that different frequencies, supported by orthogonal populations of neurons (e.g., Gloveli et al. 2005), can vary their activity independently of each other (which would be evident should theta- to gamma-power exhibit little to no correlation). Rather, these observations indicate that, across layers, gamma increases power in proportion to the amount of theta activity, supporting the hypothesis that external neural drive (theta) provides the activity that supports the amplitude of the gamma oscillation.

Although there was a significant, positive correlation in between theta and gamma power and power increases with velocity, it may be argued that these results do not fully encompass cross-frequency interactions, let alone demonstrate dependency.Therefore, we investigated the average cross-frequency phase-power coherency (Colgin et al. 2009) as a function of velocity. Across layers, gamma power-theta phase coherence increased as a function of velocity (**Fig. 5**). As theta transitions from a sinusoidal oscillation to a “saw-tooth” shape with increasing velocity, higher-order harmonics are cast in the spectral decomposition (Buzsáki et al. 1983; Sheremet et al. 2016; Terrazas et al. 2005). Notably, phase-power coherence also increased between the harmonics of theta with velocity as well as between the phase of the 16Hz harmonic and the power of gamma. As the 16Hz is not an independent oscillation from theta, but simply an oscillatory deformation, the interaction between theta and gamma encompasses both the 8hz and 16Hz component. Therefore, in order to quantify the power-phase coherence increase with velocity, the average coherence was calculated for theta phase (8Hz)-gamma power and 16Hz harmonic phase-gamma power. Calculating the difference between the high velocity bin (>35 cm/s) and low (5-15 cm/s) revealed a consistent increase in gamma power and the phase of the theta complex (**Fig. 6**).

**Figure 5:**
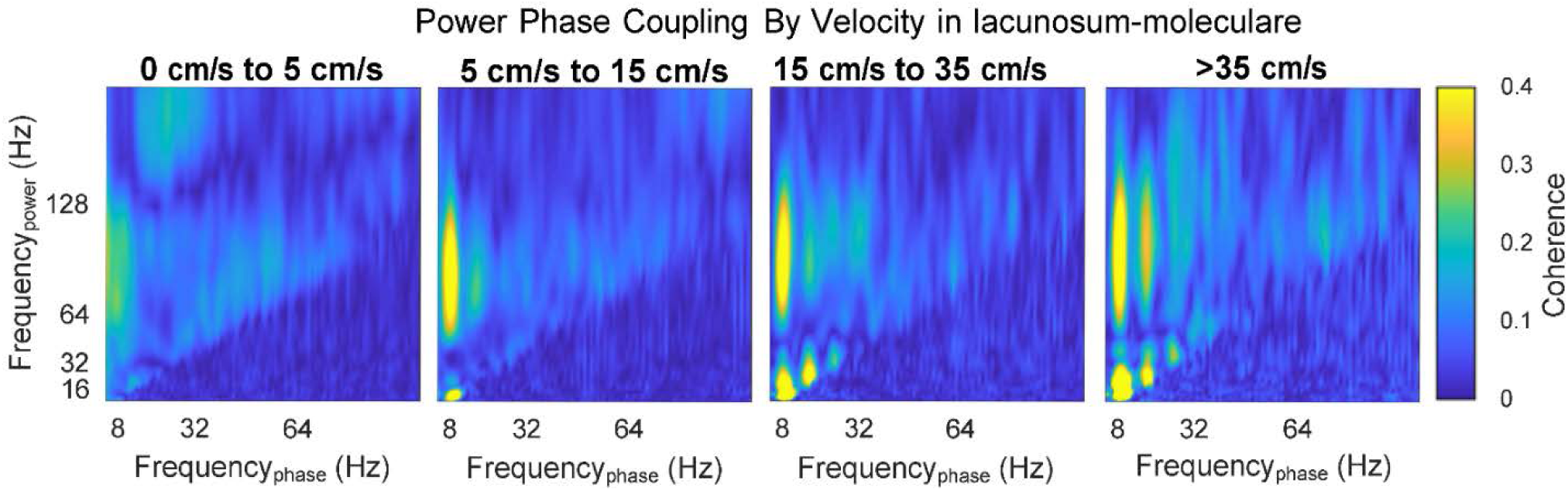
Example cross-frequency coherence between oscillatory power and oscillatory phase as a function of velocity for the lacunosum-moleculare. Note that, as velocity increases, there are low frequency interactions between theta and its harmonics (sub 40Hz). As harmonics by definition are integer, phase locked components to a fundamental oscillation, and the deformation of theta from sinusoid to sawtooth increases with velocity (Buzsáki et al. 1983; Sheremet et al. 2016; Terrazas et al. 2005), this effect is to be expected. Interestingly, there is a notable increase in the coupling between theta phase and gamma power – as well as theta harmonic phase and gamma power-with velocity.

**Figure 6:**
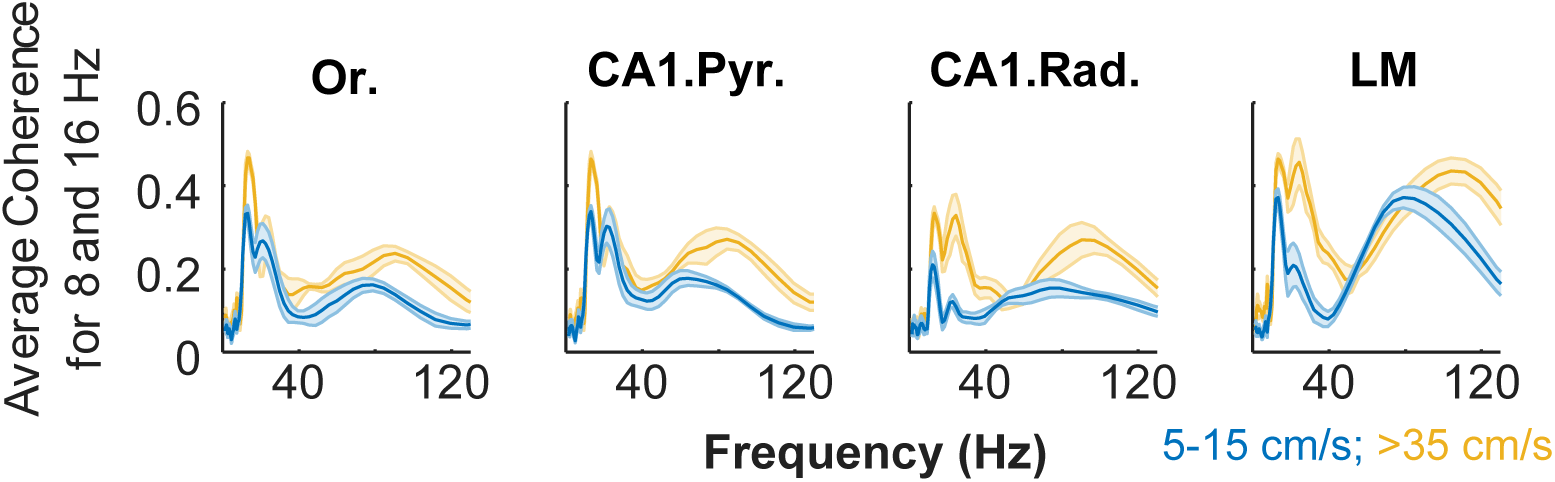
Average cross-frequency coherence for theta and the 16Hz theta harmonic phase. By averaging the coherence for these specific phases, there is a considerable change in cross-frequency coherence with velocity. Each subplot shows the individual layers of the hippocampus with yellow as the high, >35 cm/s velocity bin and blue as the 5-15 cm/s bin (Or., oriens; CA1.Pyr, pyramidal layer; CA1.Rad., radiatum; LM, lacunosum-moleculare). The shaded errorbars depict the mean and standard error of the mean (n = 5). Note that in the gamma range, greater than 50Hz, there is both an increase in coherence that is associated with a shift of the maximum value to higher frequencies (see Fig. 3). The changes in the low value frequencies, less than 40Hz, are attributed to the higher order harmonics of theta (e.g., 24 and 32Hz; see bicoherence analysis below).

As the power and phase of theta both exhibited a strong effect on gamma power, which increases in magnitude as a function of velocity, it suggests that the two oscillations are in fact part of a unitary process. Rather than the hippocampal spectra being akin to decomposition of an orchestra, with different frequencies attributable to specific, non-interacting instruments, it appears that there is a direct dependency that is analogous to ocean waves in which waves of different length form a cooperative system in which they sustain each other (e.g., Longuet-Higgins 1992). Stated simply, the LFP of the hippocampus reflects a multiscale phenomenon, driven by a flow of energy from low to high frequencies across the spectrum (Buzsaki 2006). In this energy cascade framework, it would suggest that theta-gamma interactions are dependent, with the degree of phase-phase interactions directly proportional to energy/activity in the network. However, the analytical approaches of cross-frequency phase-phase coupling have been recently called into question, due to distortions in filtering and not accounting for harmonic (Scheffer-Teixeira and Tort 2016). As described in the methods, bicoherence has been used extensively to quantify the degree of cross-frequency phase-phase coupling in the LFP, investigating the relationship between a wide band of frequencies and can be implemented to directly identify harmonics. Therefore, we investigated the degree of phase coupling within hippocampal layers as a function of velocity using bicoherence analyses.

As previously reported, bispectral estimates of theta and the respective harmonics show significant variability with rat running speed, with limited significant interactions at velocity < 10cm/s (**Fig. 7**). In contrast, at higher running speeds (i.e., velocity > 40 cm/s, **Fig. 8**) the bispectral map shows a rich phase-coupling structure. In agreement with the observations of power spectral density evolution (Fig. **3**), strong phase coupling develops between theta (theta frequency *f*_θ_ ≈ 8 Hz) and its harmonics (*f*_θ_, with n = 2,3,4, …), with peaks detectable at frequencies below ∼50 Hz, at various strengths, across all layers. It is worth emphasizing that the development of theta harmonics strongly coupled to theta indicates a nonlinear deformation of the theta rhythm toward positive skewness and asymmetry and should not be attributed to the generation of additional, statistically independent rhythms. Rather, the saw-tooth shape of theta causes a 48 Hz frequency to register in the oscillatory decomposition of LFP.

**Figure 7:**
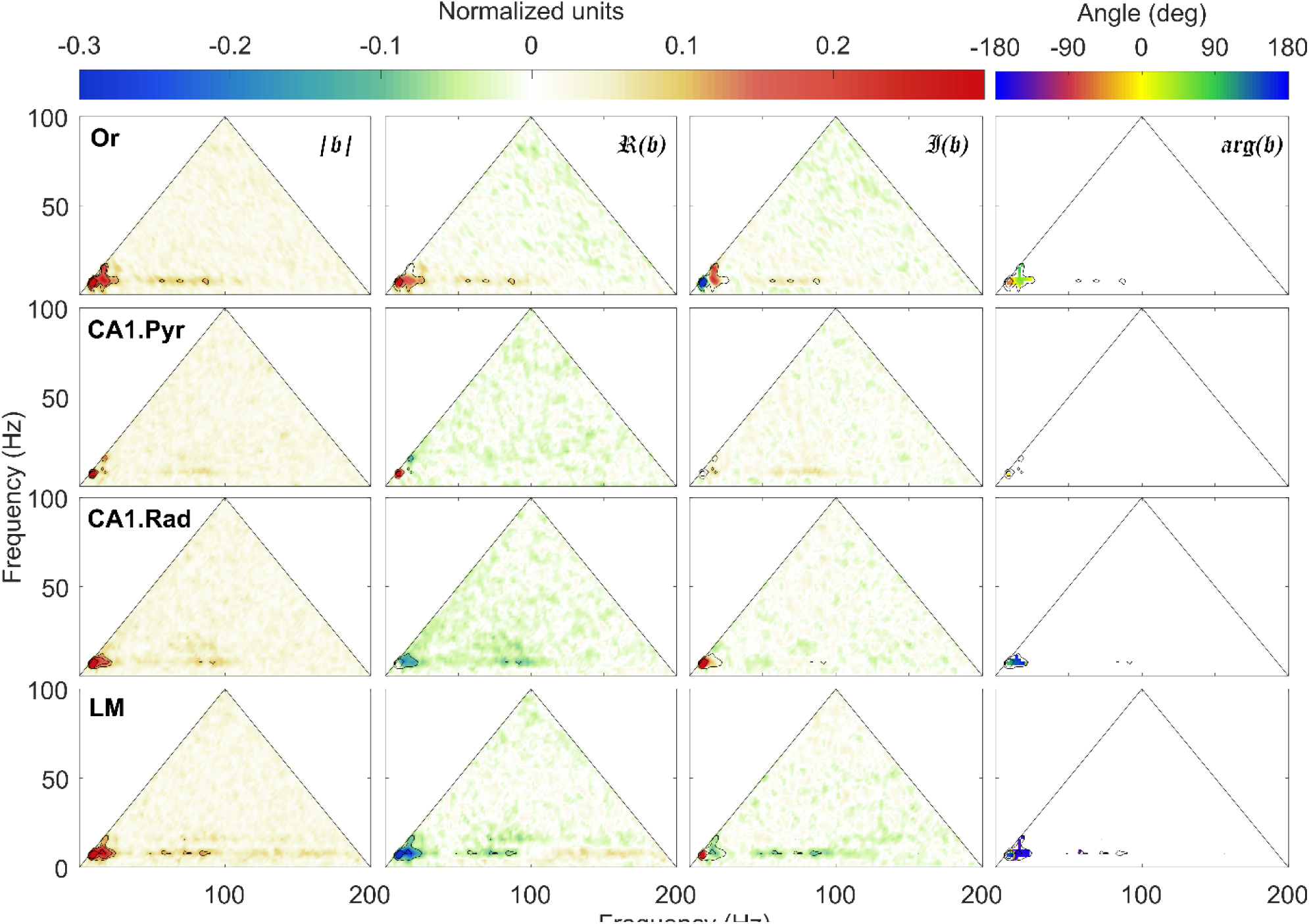
Normalized bispectrum (equation 11) at low speeds (*v* < 10 cm/s). An in-depth explanation of the bicoherence plot can be found in Sheremet et al. (2016) Briefly, the triangular region represents the area containing non-redundant information for the discrete Fourier transform. Columns (left to right) show the bicoherence |𝔟|, the real ℜ(𝔟) and imaginary 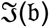 parts of the normalized bispectrum 𝔟𝔟, and the biphase arg(𝔟). The first three can be interpreted as nonlinearity strength, and measures of the contribution of different triads to the skewness and asymmetry of the LFP. Peak in the bispectral estimate represents a phase-coupled triplet (*f*_1_, *f*_2_, *f*_1_ + *f*_2_), where *f*_*j*_, j = 1,2 are frequency bands in the Fourier representation. The observations in each row correspond to a given hippocampal layer. For each layer, black contours mark the significant bicoherence value of 0.1 (with 300 DOF, zero-mean bicoherence is <0.1 at 95% confidence level.

**Figure 8.**
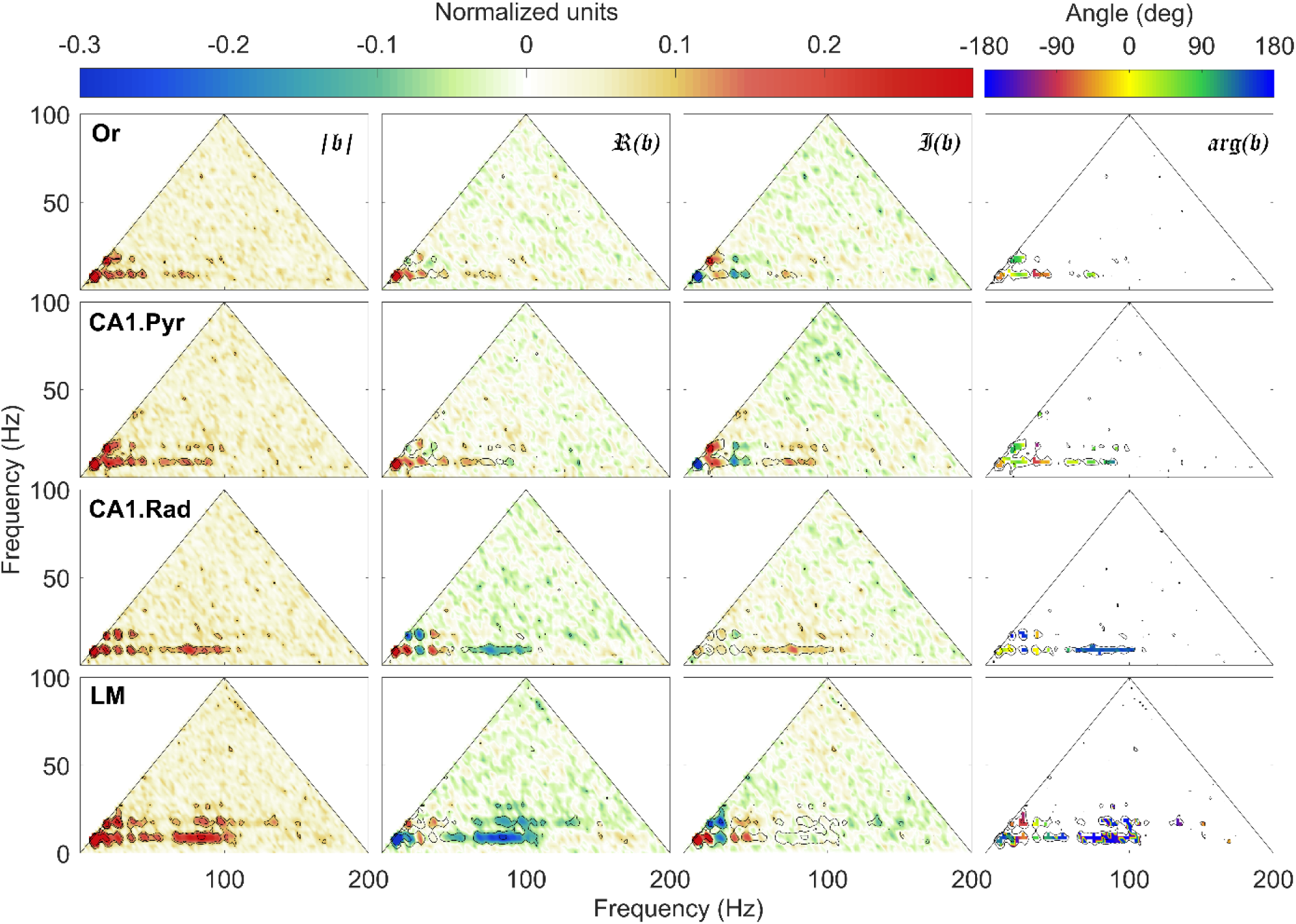
Normalized bispectrum at high speeds (*v* > 35 cm/s). Same organization as in figure 5. Note the development of multiple regions of high bicoherence |𝔟𝔟|, notably: 1) t theta (theta frequency f_θ_ ≈8 Hz) and its harmonics (nf_θ_, with n = 2,3,4, …), with peaks detectable up to 48Hz across all layers, and 2) theta and a wide gamma band spanning approximately the interval between 60 Hz to 100 Hz. The LM exhibits the richest nonlinear structures of the four layers examined with gamma coupling to the first harmonic of theta.

Furthermore, the increase of theta power and nonlinearity is accompanied by the development of significant phase coupling between theta and gamma oscillations, related to interaction between rhythms, rather than to nonlinear shape deformation.Theta-gamma coupling covers the entire gamma frequency band (the wide bands located at 50 Hz < *f*_1_ < 100 Hz and *f*_2_ ≈ 8 Hz). The effect is again strongest in the LM, where interacting triads of the form (*f*_γ_, *f* _θ_, *f* _γ_ + *f* _θ)_, and (*f* _γ_, 2 *f* _θ_, *f* _γ_ + 2 *f*) are obvious, where *f* is a frequency in the gamma band. As the location and magnitude of the bicoherence peaks identify and measure the intensity of cross-phase interactions, the strength of the nonlinear coupling can be quantified by integrating the bicoherence over the region of interest (**Fig. 9**). In agreement with the results presented in Sheremet et al. (2016), the nonlinearity was significantly greater at faster running speeds (F_[1,16]_=31.09, p = 0.001; repeated-measures). Moreover, nonlinearity also varied significantly as a function of CA1 laminae (F_[3,16]_ =11.96, p = 0.001), with the LM being significantly more nonlinear compared to the other 3 laminae (p < 0.002 for all comparisons, Tukey HSD). The bicoherence values across the Or, CA1.Pyr and CA1.Rad, however, were not significantly different (p > 0.76 for all comparisons). The difference in nonlinearity between low and high velocities did not significantly interact with layer (F_[3,16]_ = 0.21, p = 0.89), indicating that the impact of faster speeds on nonlinearity was similar across all CA1 laminae.

**Figure 9.**
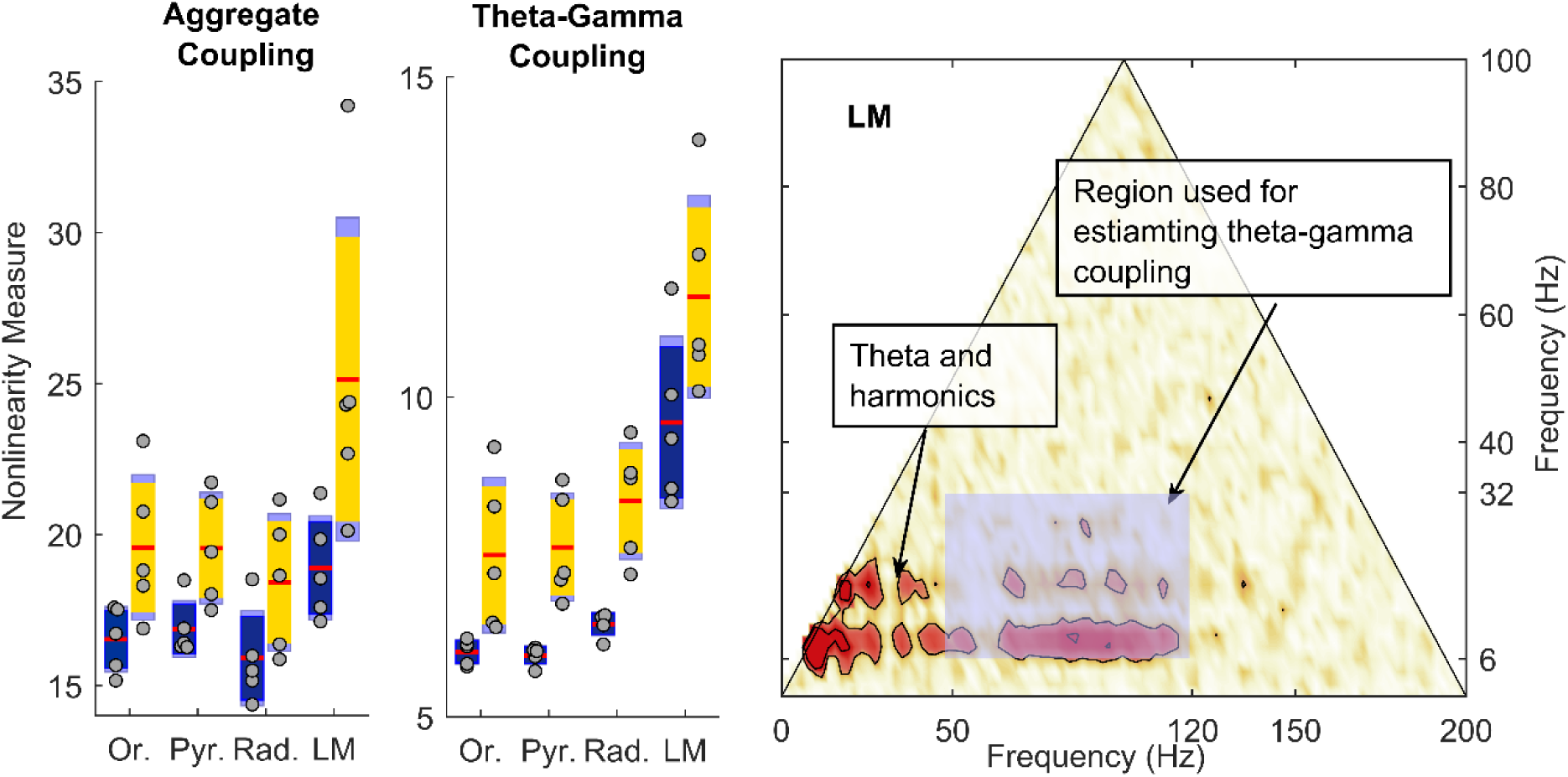
Estimate of nonlinear cross-frequency phase coupling. Left: Total overall strength of nonlinearity, estimated by summing the bicoherence over its entire definition domain. Estimates are shown for 5-15cm/s speed bin (blue) of >35cm/s speed bin (yellow). Middle: Strength of nonlinear coupling between theta (and hamonics) and gamma rhythms estimated by summing the bicoherence values in the shaded rectangle of right panel. The LM exhibits a significant larger coupling strength compared with the other layers. Both total and strictly theta-gamma coupling strength show significant variability as a function of rat speed and hippocampal layer. Note that the y-axis differs between nonlinearity measures in order to optimize for display of the theta-gamma-gamma nonlinearity. Red is the mean, solid dark blue or yellow is 1.96 of the standard error of the mean (S.E.M.) and light blue denotes the boundaries of 1 standard deviation. Right: Region used for estimating the strength of theta-gamma coupling is a rectangle covering the intervals [6 Hz, 32 Hz] for theta and [50 Hz, 120 Hz] for gamma.

When the quantification of nonlinearity was restricted to the theta (6-32 Hz) and gamma (50-120 Hz) frequency ranges (Fig 7), there was also a strong effect of velocity on nonlinearity (F_[1,16]_ =38.53, p = 0.0001; repeated measures). Within the theta and gamma frequency ranges there was also a significant effect of layer on nonlinearity (F_[3,16]_ =19.87, p = 0.001), that did not significantly interact with velocity (F_[3,16]_ = 0.48, p = 0.70). Interestingly, the theta-gamma bicoherence values in the LM were significantly greater than the other 3 laminae (p < 0.001 for all comparisons; Tukey HSD). The nonlinearity between the Or, CA1.Py and CA1.Rad was not significantly different, however (p > 0.32 for all comparisons; Tukey HSD).

## Discussion

The current paper reports the novel finding of power-power, phase-power, and phase-phase coupling between theta and gamma oscillations that is dependent on velocity, which varies in a layer dependent manner. Macroscopically, bouts of input at theta frequency provide the excitation/inhibition to organize synaptic activity rhythmically. Neurons within the entorhinal-hippocampal circuit are tuned to resonate at theta frequencies, evident in the ability of the hippocampus to locally generate theta when provided sufficient input (Bland et al. 1988; Fraser and MacVicar 1991; Kamondi et al. 1998; Kocsis et al. 1999; Konopacki et al. 1987). Therefore, when strongly driven, local circuits find a temporally stable interaction. This drive, in turn, activates small circuits, which – when “kicked hard”- provide a mechanism for endogenous timing. For example, this theta paced drive could conceivably cause basket cells to fire gamma frequency bursts at theta frequency (Bragin et al. 1995; Ylinen et al. 1995b). The consequence of this organization is that neuronal activity of both pyramidal cells and interneurons are organized to both the relatively slow theta oscillation and the faster gamma oscillation (Buzsáki and Chrobak 1995).

Increasing the total amount of activity into a network will have multiple consequences including more coincident synaptic events (reflected as increases in local field potential power). There is a dramatic increase in firing rates across hippocampal neurons with velocity (Hirase et al. 1999; Maurer et al. 2005; McNaughton et al. 1983). Moreover, running speed alters hippocampal sequence compression of place cell spikes (Maurer et al. 2012). Because the single units change their dynamics, and the LFP is generated by membrane events, it stands to reason that there must be associated changes in LFP power and gamma frequency (Ahmed and Mehta 2012; Chen et al. 2011; Kemere et al. 2013; Zheng et al. 2015). It is worth noting that there is a low fraction of observed gamma locking between CA3 or entorhinal LFP and CA1 pyramidal neurons (Schomburg et al. 2014). These observations support the idea that gamma can provide a measure of upstream temporal dynamics but fails to entrain the individual units (Buzsáki and Schomburg 2015). Rather, as the distal dendrites operate as a low-pass filter (Golding et al. 2005), theta-gamma paced synaptic input in the lacunosum-moleculare or radiatum are reverted to just theta at the soma (Vaidya and Johnston 2013). Nevertheless, the lacunosum-moleculare exhibited the highest theta-gamma coupling as a function of velocity suggesting that theta-modulated gamma inputs are conveyed from the entorhinal cortex (Fernández-Ruiz et al. 2017), which exhibits a high amount of sensitivity to path-integration and self-motion (Kropff et al. 2015; Moser and Moser 2008; Sargolini et al. 2006).

The interplay between theta and gamma rhythms as a function of velocity is not to suggest that theta is a required prerequisite for gamma. There are two considerations with respect to this point. First, the power spectra density of the local field potential demonstrates and ∼ 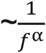 slope, indicating that any frequency has the potential of being isolated. Power will exist across all frequencies, with low frequencies exhibiting the highest amplitude. Therefore, while there may not be a large peak in theta power, there still may be enough activity in the network that leads to local circuit entrainment at gamma frequency. Experimentally, this has been observed in situations in which severing the fimbria fornix or lesioning the entorhinal cortex severely diminished theta power, but yielded increases in gamma power (Bragin et al. 1995; Buzsáki et al. 1987). This highlights the second consideration, that it is the characteristics of neurons and the local network that defines the ability to capture the incoming activity. In the absence of entorhinal input, it is tenable that the network undergoes reorganization to compensate for the loss of input. This compensation results in an easier entrainment of local circuits (gamma) that occurs with weak, low frequency input across the population. Other aspects theoretically modulate the ability of local circuits to capture the global input including age-related synaptic changes and pharmacological manipulation.

The current manuscript links prior findings of theta-gamma interaction in the service of cognition (e.g., Jensen and Colgin 2007; Lisman and Jensen 2013; Montgomery et al. 2008; Tort et al. 2009; Tort et al. 2008) to other studies that have observed layer dependent theta-gamma effects during exploration (Belluscio et al. 2012; Lasztóczi and Klausberger 2016; or using datasets with and with out a mneumonic requirement Schomburg et al. 2014). The current data, however, point to a new perspective. In opposition to the idea that gamma and theta are driven by minimally overlapping mechanisms that couple under certain behavior conditions, we propose that coupling is enhanced in situations in which there is increased input into the network. Under increased input into the network, which occurs at faster running speeds or instances of high cognitive load, theta-gamma coupling could be utilized by local circuits to support higher-cognitive processes. For example, over the course of learning, theta paced inputs may result in higher power gamma oscillations through experience-dependent plasticity. In this manner, in the awake-behaving animal, theta and gamma are interdependent, coupled by the cascade of energy across spatial and temporal (frequency) scales (Buzsaki, 2006).

This perspective, however, runs counter to the recent theory of oscillatory multiplexing in the brain, where individual frequencies carry different components of information (e.g., Akam and Kullmann 2014; McLelland and VanRullen 2016). For example, it has been suggested that communication across brain regions may occur in a frequency specific manner and aspects of cognition can be determined through calculating cross-regional coherence, with different bands conveying different psychological components (Knight and Eichenbaum 2013; Siegel et al. 2012; Watrous et al. 2013). Moreover, neuron populations with heterogeneity in the preferred phase coupling frequency would theoretically provide a mechanism to communicate and organize the spike firing times of multiple, distant cell assemblies in a frequency specific manner (Canolty et al. 2010). However, this entrainment by oscillatory coherence may be an over-simplification. For example, the dendrites of hippocampal pyramidal neurons operate as a low pass filter (Golding et al. 2005; Vaidya and Johnston 2013), which would suggest that gamma frequency activity within the lacunosum-moleculare (the afferent input layer of the entorhinal cortex) has little consequence on the gamma frequency activity within the pyramidal layer. This is evident in the relatively low coherence of gamma frequences between the medial entorhinal cortex and pyramidal layer when compared to theta coherence between the same two regions (∼0.3 and ∼0.85 respectively, Colgin et al.2009). Therefore, rather than coherence, external inputs entrained at gamma may instead provide the necessary input to coordinate the phase of theta within a network (Lopez-Madrona et al. 2018). Tersely, entorhinal input at gamma frequency supports theta paced dynamics, and the energy cascade begins again. This line of reasoning raises the critical consideration that marginal coherence values (stated differently, incoherence) may not be biologically meaningful and coupling measures may risk interchanging correlation for consequence. Rather, it may not be the frequency of communication that is important, but the total amount of activity into the network.

The cascade theory suggests that it is improvident to assign a particular psychological process to a specific frequency for the reason that, in the awake behaving animal, the gamma rhythm is shaped by the activity in the theta band making the two oscillations interdependent. Indisputably, the iterative processing and nonlinearity of neural circuits make it implausible to assign a cause and effect relationship between a cognitive process and a narrow oscillatory band. The local field potential, however, is diagnostic of the underlying interactions, providing information to the experimenter with respect to *how the network is predisposed to organize*. In fact, synaptic events – contributing the ionic flux that shapes the LFP – have no knowledge with respect to whether they are supporting “gamma” or “theta.” Parsing oscillatory frequencies to specific biophysical mechanisms, circuits or behavioral correlations for the purposes of higher-cognition loses relevance if the nervous system does not make the distinction (unless it is done for diagnostic purposes as described in the condition).

While assigning certain frequency oscillations may be gratifying to the experimentalist, Fourier decomposition is an analytical tool, far removed from synaptic or circuit dynamics. For example, often it is tempting to construct a power spectra and ask a question regarding deviations away from the 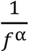 slope (e.g., after whitening). In the current manuscript, we demonstrate that the spectral slope *α* is not constant throughout the entire frequency range, demonstrating a methodological flaw to standard whitening techniques. Moreover, our spectral decomposition revealed 16, 24, 32, 40 Hz and higher harmonics of theta (Sheremet et al. 2016). While the decomposition “uncovers” these oscillations, the harmonics are the consequence of a single aggregate process in which theta changes from a sinusoid into a sawtooth with high asymmetry and skewness. Furthermore, there is an associated loss of power between ∼1-6 Hz as running speed increases (Fig. 3). Interestingly, the loss of lower frequency power along with the development of the harmonics suggests that, as the amount of input increases, the initial effect will not necessarily be the entrainment of local circuits at gamma frequency, but increasing the network activity in the theta band This hypothesis is supported by the observation of inhibition dependent theta resonance (Stark et al. 2013). When there is little to no input, interneurons that resonant at theta frequency will express incoherent entrainment between 1-6 Hz. As input increases, these neuron begin to resonate, with power redistributing into theta. Finally, as hippocampal activity increases with velocity, the oscillatory energy begins to extend beyond the theta resonance and begins to entrain local circuits at gamma frequency.

This theory of energy cascade is not novel. In fact, it shares similarity with the concepts of self-organized criticality (Bak et al. 1987). Self-organized criticality was broadly used to describe any system in which there is an inverse correlation between oscillatory power and oscillatory frequency. This concept later found an analogous home in neuroscience, describing both the shape of the power spectra as well as neuronal avalanches (Beggs and Plenz 2003; Beggs and Timme 2012; Plenz and Thiagarajan 2007). With these insights, it is possible to consider the mechanisms by which the brain gives rise to cognition by relating activity through neural circuits.

## Acknowledgements

This work was supported by the McKnight Brain Research Foundation, and NIH grants-Grant Sponsor: National Institute on Mental Health; Grant Number: R01MH109548 and a Diversity Supplement to NIH grant R01MH109548 (JPK). We would like to thank Drs. Gyuri Buzsaki, Kamran Diba and Eric Schomburg for comments on an earlier draft of the manuscript.

